# A transcriptional regulator gene *TRG1* of *Trichoderma longibrachiatum* is involved in the regulation of peptaibols synthesis

**DOI:** 10.64898/2026.01.07.698138

**Authors:** Weishe Hu, Aizhi Ren, Mengjiao Guan, Xiaoting Wang, Ming Li, Xiusheng Zhang, Peibao Zhao

## Abstract

*Trichoderma* species are recognized for their robust biocontrol capacity, primarily due to the production of peptaibols, potent secondary metabolites that exhibit antibacterial properties, induce cell apoptosis, and enhance plant disease resistance. To elucidate the regulatory mechanism of peptaibol synthesis, we investigated the role of a transcription regulator gene *TRG*1 (designated as *TLTRG*1), which we identified within a non-ribosomal peptide synthetase (NRPS) gene cluster of *T. longibrachiatum* SMF2. We successfully created a *ΔTRG*1 gene knockout mutantusing homologous recombination. Phenotypic analysis indicated that the *ΔTRG*1 mutant maintained wild-type growth rate, colony morphology, and spore production. However, *ΔTRG*1 exhibited a significantly reduced inhibition rate against the plant pathogen *Botrytis cinerea* and a lower efficacy in controlling gray mold on detached rose leaves. High-performance liquid chromatography (HPLC) analysis provided direct molecular evidence, showing that peptaibol production in the mutant was 2.5 times lower than that of the wild-type strain. These findings conclusively establish that TRG1 functions as a key positive transcriptional regulator essential for high-yield peptaibol biosynthesis by modulating NRPS gene expression. This gene represents a critical molecular target for bioengineering strategies aimed at enhancing the biocontrol efficacy of *Trichoderma* strains.

**The Importance section:** *Trichoderma* species are known to produce peptaibols, which exhibit antibacterial properties, induce cell apoptosis, and enhance plant disease resistance. However, the mechanisms underlying the biosynthesis and regulation of peptaibols remain unclear. In this study, we investigated the role of a transcription regulator gene located within a non-ribosomal peptide synthetase (NRPS) gene cluster of T. longibrachiatum SMF2, which we designated as TLTRG1 (TRG1).

Phenotypic analysis indicated the inhibition rate of ΔTRG1 against Botrytis cinerea and its effectiveness in controlling gray mold were significantly reduced compared to the WT strain. High-performance liquid chromatography (HPLC) analysis revealed that the production of peptaibols in the mutant was significantly decreased. These findings suggest that the TRG1 gene may play a crucial role in regulating the expression of NRPS genes, thereby affecting the biosynthesis of peptaibols.

Therefore, it can be concluded that the gene TRG1 is a molecular target for bioengineering strains with enhanced biocontrol efficacy.

## 1. Introduction

As global living standards rise, there is an intensifying focus on health, driving a growing interest in environmentally friendly agents for the prevention and control of crop diseases (Chang et al., 2024). *Trichoderma* is the most commonly used fungus, widely distributed in soil, and plant rhizospheres (Ramirez et al., 2019; Guzmán et al., 2024). Its primary biocontrol mechanisms include competition, antibiosis, and the induction of host resistance (Papavizas, 1985; Howell, 2003). However, the widespread application of *Trichoderma* is limited by the inherent drawbacks of biopesticides, such as slow efficacy, narrow control spectrum, and susceptibility to environmental conditions (Woo et al., 2023; Schmaltz et al., 2023). To overcome these limitations and expand its agricultural utility, a feasible strategy is to investigate the biocontrol mechanisms of *Trichoderma* at a molecular level, specifically by developing efficient biocontrol agents from its secondary metabolites. *Trichoderma* produces various compounds, including peptaibols, which exhibit important functions such as antibacterial activity, inducing cell apoptosis and enhancing plant disease resistance. These peptaibols are synthesized via non-ribosomal peptide synthetases (NRPSs) pathways (Song et al., 2006; Luo et al., 2010; Shi et al., 2012; Zhao et al., 2018). In recent years, extensive research has elucidated the structure, function and biosynthetic pathway of NRPSs and their peptaibols. (Grünewald and Marahiel, 2006; Degenkolb et al., 2008; Madsen et al., 2016; Matt et al., 2019).Despite their molecular understanding, the broad agricultural and pharmaceutical application of NRPS-derived peptaibols faces significant challenges. NRPSs are the largest enzymes discovered to date, making artificial expression difficult. Furthermore, current genetic engineering techniques are typically limited to NRPSs that produce small peptides (3-5 amino acids) (Degenkolb et al., 2008; Miyanaga et al., 2019; Matt et al., 2019).Therefore, strengthening the basic research related to the synthetic mechanism of peptaibols is crucial. Specifically, investing the regulatory mechanisms of peptaibol synthesis at the transcriptional level and seeking to improve th peptaibol yield by enhancing NRPS expression through bioengineering is essential to unlock the full potential of *Trichoderma* and peptaibols in controlling of plant diseases.

In this study, we aimed to identify and characterize a transcriptional regulator involved in peptaibol synthesis. We first used bioinformatics analysis to identify a transcriptional regulator (*TRG*1) within an NRPS gene cluster in *T. longibrachiatum* SMF2. We then created gene knockout mutants and demonstrated that the antibacterial activity and production of peptaibols in the *ΔTRG1* mutant were significantly reduced. These findings strongly indicate that *TRG*1 is a key transcriptional regulator of peptaibol synthesis in *T. longibrachiatum* SMF2 and establish a new target for the bioengineering-driven improvement of biocontrol agent production.

## 2 Materials and methods

### 2.1 Fungi

The wild type strain *Trichoderma longibrachiatum* SMF2 was obtained from the Key Laboratory of Microbial Technology at Shandong University. The plant pathogen, *Botrytis cinerea* T_4_ was obtained from Prof. Dickman Marty of Texas A&M University. Both *T. longibrachiatum* SMF2 and *B. cinerea* T_4_ were routinely cultured on potato dextrose agar (PDA) or potato dextrose broth (PDB) medium at 25 ± 2°C (Zhao et al., 2018). Mycelium was collected and genomic DNA was extracted using a fungal genomic DNA kit (Invitrogen) for use as a template in subsequent PCR amplification and for molecular validation experiments..

### 2.2 Bioinformatics analysis of NRPS Claster

Bioinformatics analysis of the genomic sequence surrounding the NRPS gene *tpx1* was conducted to identify the complete NRPS cluster and associated genes. The coding sequences and introns within the cluster were identified using the software DNAman and MEGA 11. The promoter and homologous genes were analyzed using online software promoter 2.0 and BLASTn to infer the functions of the identified genes, particularly potential regulators (like transcription factors) or tailoring enzymes involved in peptaibol synthesis.

### 2.3 Construction of knockout vectors of the gene *TRG1* and mutant screening

Based on the gene homologous recombination theory, the *TRG1* gene-disruption vector pUCATPH-*TRG1* was constructed. This vectorcontains a hygromycin B resistance gene *hph* as a selectable marker, which was sourced from the pUCATPH plasmid,acquired from the Key Laboratory of Agricultural Microbiology in Shandong Province, China. DNA of *T. longibrachiatum* SMF2 was isolated and used as s a template to PCR amplify the upstream and downstream fragments of the *TRG1* gene using primer pairs LG1-1/LG1-2 and LG1-3/LG1-4 respectively, followed by primer pairs LG1-1P/LG1-2P and LG1-3P/LG1-4P, harboring appropriate restriction sites. The resulting fragments were recovered and purified for use as upstream and downstream recombination arms in the construction of the mutant vector. All the primers used in the experiment were shown in Table 1.

**Table 1.**
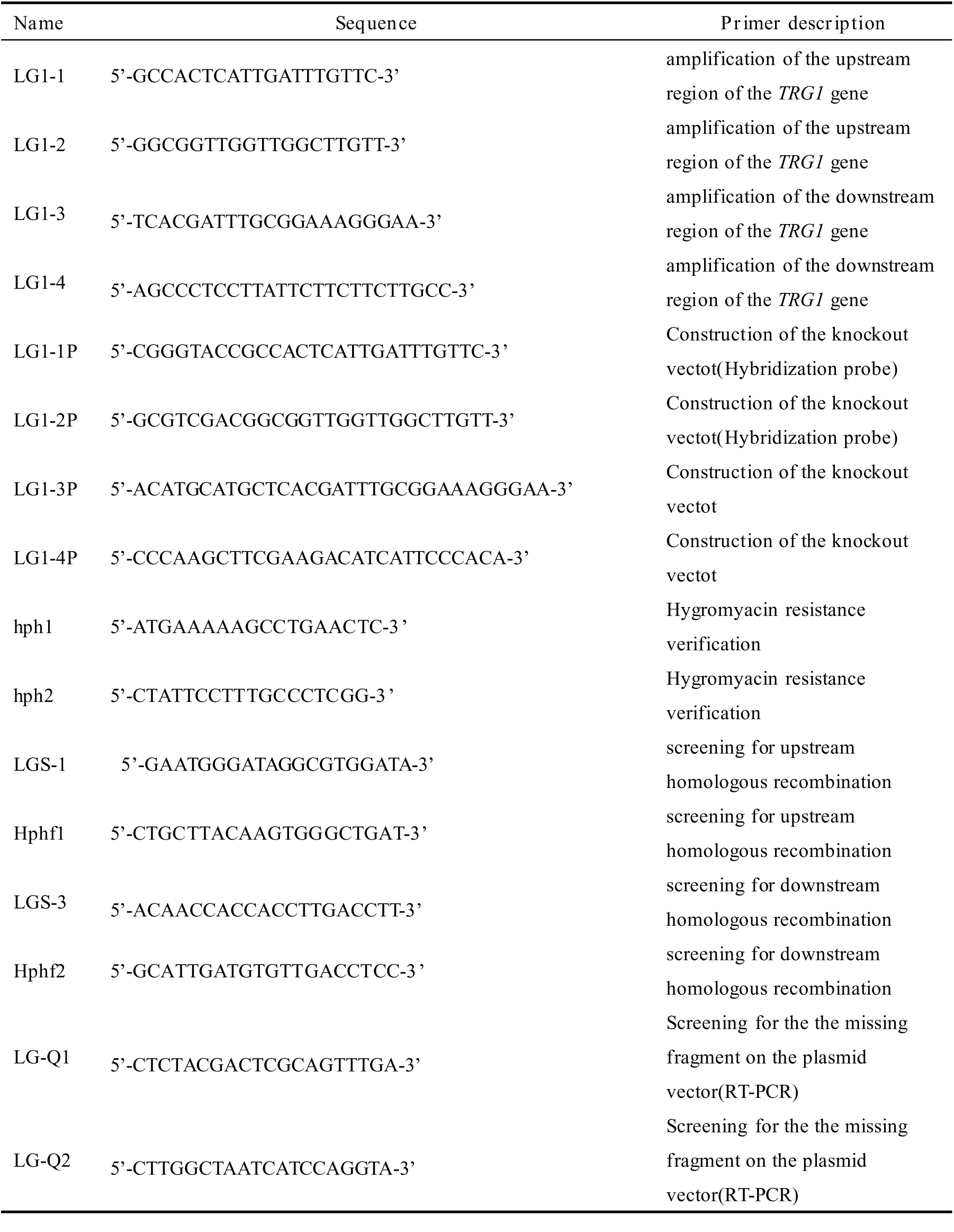
List of primers used in the expriment.

First, the fragment TRG1-U, which was amplified using the primer pair LG1-1P/LG1-2P, and the vector pUCATPH were digested with Sph I and Hin dIII, and then ligated by T4 ligase. Then, another DNA fragment TRG1-D, which was amplified by the primer pair LG1-3P/LG1-4P, was digested with Kpn I and Sal I and ligated into the recombinant plasmid pUCATPH-U, which was digested using the same enzymes. The recombinant vector pUCATPH-TRG1 was identified and subsequently used to transform T. longibrachiatum SMF2 protoplasts utilizing a restriction enzyme-mediated integration (REMI) transformation technique (Yechun et al., 2007; Zhao et al., 2010). Transformants were selected on hygromycin-containing medium. Positive transformants were initially verified by PCR analysis.Then Southern hybridization and RT-PCR were performed to confirm the single-copy integration of the hph cassette and the absence of TRG1 gene transcription in the knockout mutant ΔTRG1. All molecular experiments followed standard protocols (Jeo et al., 2001).

### 2.4 Phenotypic characterization of the mutant *ΔTRG1*

The wild-type strain (WT) and the mutant *ΔTRG1* were cultured on PDA medium for 7 days to assess growth rate, colony morphology, spore production, and other phenotypic characteristics. Assays were conducted in triplicate (n = 3). The colonies diameter of the WT and *ΔTRG1* were measured daily to evaluate their relative growth rate.

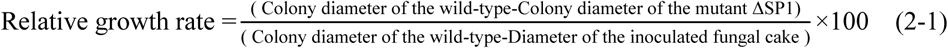

### 2.5 Biological control assays of the mutant *ΔTRG1*

The confrontation culture method was used to determine the inhibitory effect of the WT strain and mutant *ΔTRG1* on the growth of *B*. *cinerea* T_4_. The strain of the WT and mutant *ΔTRG1* were co-inoculated with *B*. *cinerea* on PDA medium. *B*. *cinerea* cultured alone was used as a negative control. Assays were conducted in triplicate (n = 3). The diameter of *B*. *cinerea* colonies were measured on day 6 after inoculation in the different treatment groups and the inhibition rate was calculated using the following formula:

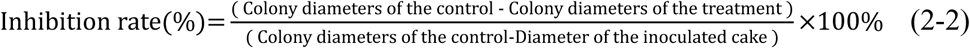

Detached rose leaves were used to evaluate the protective efficacy of the mutants against rose gray mold disease caused by *B. cinerea*. Firstly, the detached leaves were inoculated with a mycelial cake of *B. cinerea* T_4_, and placed on the filter paper spread in the culture dish, maintaining humidity with sterile water. Two days post-inoculation, the spore suspensions (1×10^6^spores/mL) of the WT and mutant *ΔTRG1* were sprayed on the leaves using sterile syringes. The control group received an equivalent volume of sterile water sprayed onto the pathogen-inoculated leaves. Assays were conducted in triplicate (n = 3). Subsequently, the disease severity of leaves was assessed on day 7 after spraying with spore suspensions of the WT and*ΔTRG1*, the disease index and protective effect were calculated according to formulas 2-3 and 2-4 respectively, to determine the control effect of the WT and*ΔTRG1* on rose gray mold disease (Zhao et al., 2018). (Disease grading criteria: 0: No symptoms;1: Lesion area <12.5% of leaf area; 2: 1.5%≤ Lesion area <25% of leaf area; 3: 25%≤ Lesion area <50% of leaf area; 4: Lesion area ≥50% of leaf area)(Zhao et al., 2018).

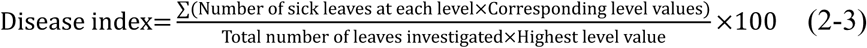

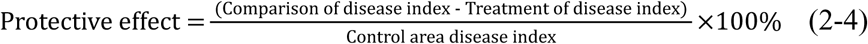

### 2.6 Antibacterial assay of the cell-free fermentation supernatant

After culturing on PDA medium for 5 days, the spores of the WT strain and the mutant *ΔTRG1* were collected and spore suspension prepared was prepared (1×10^8^ spores/mL). This suspension was inoculated into PDB medium and cultured in a shaking incubator, at 150 rpm, 25°C for five days. The culture mixture was filtered through four layers of gauze to remove hyphae, and the filtrate was centrifuged for 10 min, at 12,000 rpm. The resulting supernatant was filtered through a 0.22 µm sterile filter membrane. This cell-free supernatant was then mixed with sterile PDA in a ratio of 1:10 (v/v), mixed evenly, and poured into Petri plates. *B. cinerea* T_4_ was inoculated onto the plate and cultured at 25°C for 4 days. The diameter of the colony was measured, and the inhibition rate was calculated according to Formula 2-2.

### 2.7 Detection of induced resistance-related enzymes

The fermentation broth of the WT and mutant *ΔTRG1* was prepared as described in Section 2.6. This broth sprayed onto detached rose leaves, with sterile water was used as a comparative control. Assays were conducted in triplicate (n=3). The leaves were sampled on the 1st, 3rd, and 5th days post-treatment to measure the activities of key resistance-related enzymes: soluble protein, superoxide dismutase (SOD) and catalase (CAT). Enzyme activities were measured following the methods described in Güneş (2019).

### 2.8 High performance liquid chromatography (HPLC) analysis

The WT and Δ*TRG1* strains were cultured on PDA plates for 7 d, spores were collected, inoculated into solid fermentation medium (9g dry wheat bran with 45% moisture content, Grass powder 1g, KH2PO4 0.048 g, (NH4)2SO4 0.048 g, CaCl2 0.02 g, MgSO4•7H2O 0.02 g) for 7 d, at 28 ℃. The fermented material was extracted with 100 mL anhydrous ethanol for 4 hours. After filtration and centrifugation at 12,000 rpm, the supernatant was concentrated and drief using a rotary evaporator. The dried extract was resolubilized in 10 ml methanol and filtered through a 0.22 μm filter before HPLC analysis (Song, XY et al. 2006), The HPLC parameters were set as follows: mobile phase was methanol and water(81:19); flow rate was 1mL/min. High-performance liquid chromatography (HPLC), equipping with an ultrasonic degasser (DGU-20A5R), a binary high-pressure gradient pump (LC-30AD), an automatic sampler (SIL-30AC), a column oven (CTO-20AC), a photodiode array detector (SPD-M30A), and a communications bus module (CBM-20A). The column specifications were 250 × 4.6 mm in size, with a 5 μm C18 packing material. And the detection wavelength of HPLC is 254 nm.

### 2.9 Statistical analysis

The experimental data were analysed by ANOVA using IBM SPSS Statistics 26 and the means of different treatments were compared by using Least Significant Difference (LSD) method. Figures were plotted using OriginPro 2023(Qi et al., 2024).

## 3 Results

### 3.1 The transcription regulator gene *TRG1* is located on the NRPS gene cluster containing the NRPS gene *tpx1*

Through bioinformatics analysis, more than ten NRPS-PKS gene clusters were identified in the genome of *T. longibrachiatum* SMF2. Previous studies have shown that Trichokonin Ⅵ a 20-amino acid active peptide, is the most important active peptide produced by *T. longibrachiatum* SMF2, and its synthesis is governed by the NRPS gene *tlx1*(Xie et al., 2015; Zhou et al., 2019). We investigated the gene cluster where the NRPS gene *tlx1* is located, and found that the predicted gene cluster contains genes that are likely related to peptide modification, transport, and synthesis regulation (Fig. 1). In particular, a predicted C6 transcription factor gene (KB290*LG1*_475) and a transcription regulator gene (*TRG1*) were found, both of which were associated with gene expression regulation (Fig. 1).

**Figure 1.**
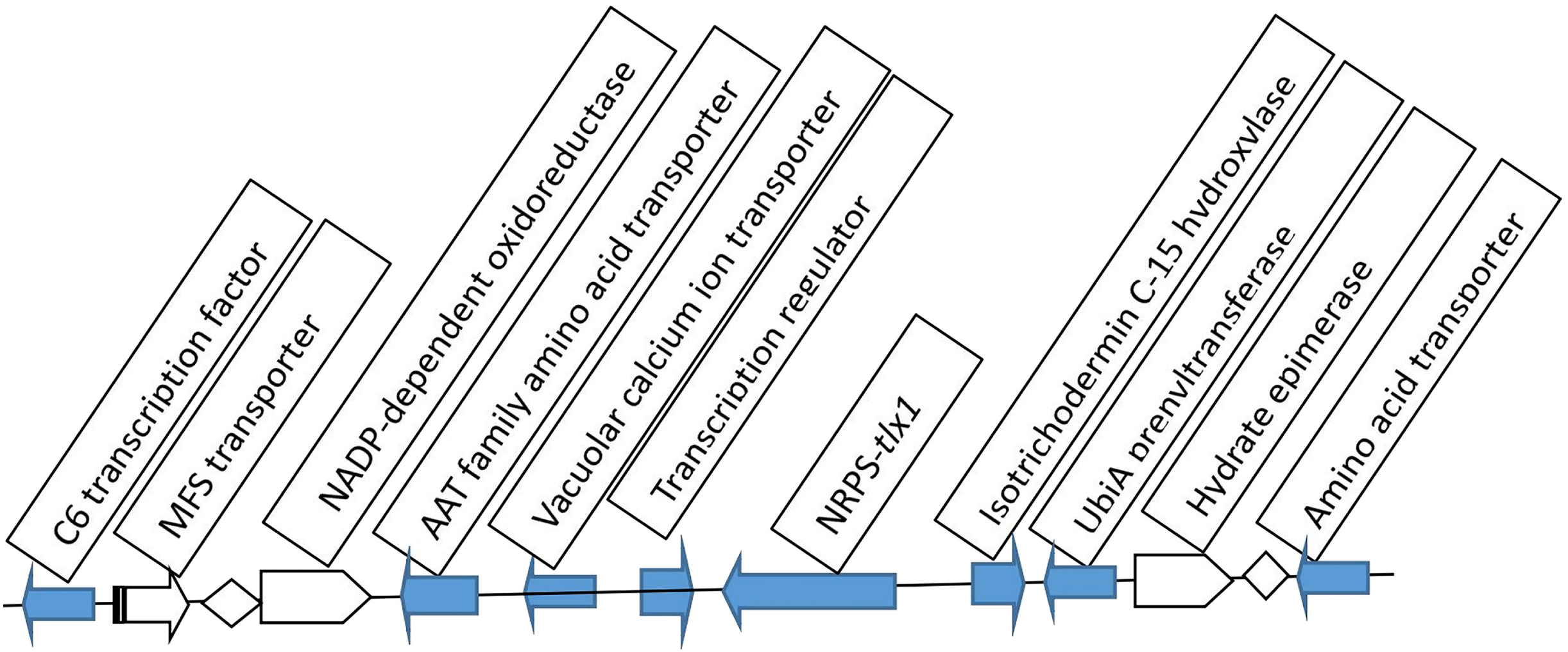
The location of the *TRG*1 gene within the non-ribosomal peptide synthetase(NRPS) gene cluster containing *tlx*1.

### 3.2 Construction and identification of *TRG1* mutant in *T. longibrachiatum* SMF2

The upstream and downstream recombination arms of *TRG1* were amplified by PCR with primer pairs LG1-1P/LG1-2P and LG1-3P/LG1-4P, yielding products of 750 bp and 1000 bp, respectively. These two fragments were inserted into the recombinant plasmid pUCATPH-*TRG1* using a two-step method, with the plasmid containing a hygromycin resistance gene *HPH*. A schematic of the construction of the recombinant vector pUCATPH-*TRG1* is presented in the Fig. 2.

**Figure 2.**
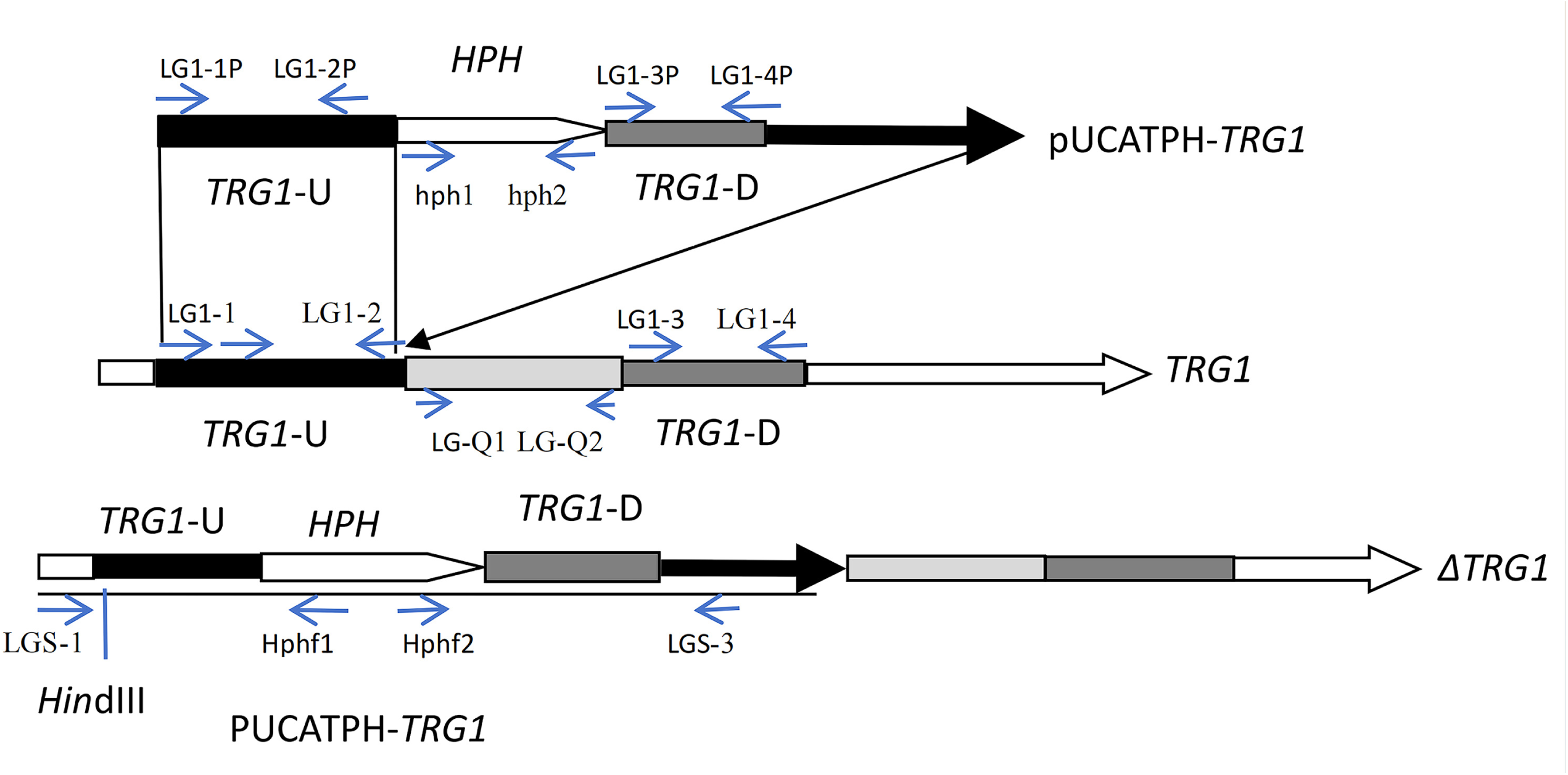
Schematic diagram illustrating the*ΔTRG*1 knockout strategy, showing the recombinant vector (pUCATPH-*TRG*1) and the predicted homologous recombination event. pUCATPH-*TRG1*: The knockout vector; *TRG1*: The *TRG1* gene; *ΔTRG1*: Diagrammatic representation of recombinant gene (The figure contains the names and positions of the gene fragments, primers, and enzyme cutting sites).

A total of 78 transformants obtained through REMI transformation were screened. After culturing for multiple generations to ensure stable inheritance of hygromycin resistance, a recombinant mutant was identified, designated as Δ*TRG1,* was identified via PCR and hybridization screening. Initial PCR and sequencing analysis confirmed the correct upstream recombination, yielding a product that contained the genomic fragment upstream of the recombination site, the upstream recombination arm, and a partial fragment of the HPH gene (Table 2). Meanwhile, the PCR results showed that the missing gene fragments of *TRG1* on the vector could be obtained from the mutant DNA(Fig. 3A). Further validation was performed by Southern blotting. When hybridization was performed with the upstream recombination arm as a probe, only one hybridization band was obtained in both the WT and mutant *ΔTRG1*(Fig. 3B), indicating a successful homologous recombination in the mutant *ΔTRG1*. If the recombinant vector was a heterologous insertion without gene recombination event, there will be two hybridization bands, one with the same as the WT (Shorter than 9,000 bp), and the other one will be longer than the recombinant vector (Fig. 2; Fig. 3A/3B). In short, the vector pUCATPH-*TRG1* was introduced into the *T. longibrachiatum* SMF2 genome via a single crossover event, resulting in the disruption of the integrity gene, without any gene fragment deletion or heterologous insertion events in the genome. A diagram of the presumed recombination model is shown in Fig. 2 and the detected recombinant sequence is shown in Table 2.

**Figure 3.**
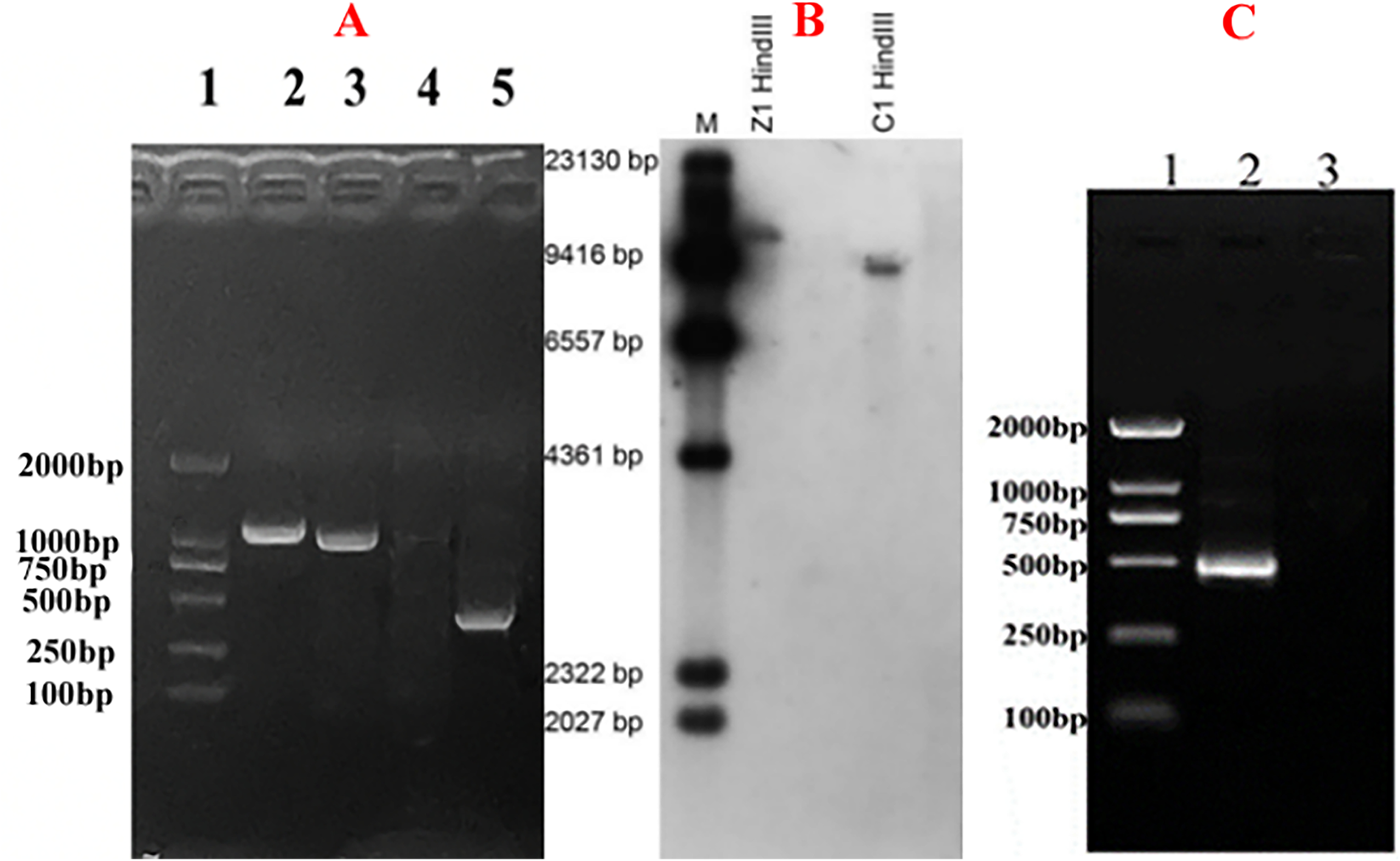
Validation of the gene recombinant in Δ*TRG1*. A. PCR analysis of the gene deletion in the mutant Δ*TRG1*. (Lane 1: Size Markers; 2: PCR amplification product using primer pair hph1/hph2; 3: PCR amplification product using the upstream recombination primers; 4: Absence of PCR product of using the downstream recombination primers; 5: PCR amplification of the missing fragment) B. Southern blotting verification of vector insertion and gene recombination in Δ*TRG1* mutant (Lane 1: Size Markers; 2. Z1:Hybridization product using Mutant Δ*TRG1* DNA digested with HindⅢ as the template; 3. C1: Hybridization product using the WT DNA digested with HindⅢ as the template). C. RT-PCR product using *TRG1* primer set on cDNA derived from WT and Δ*TRG1* strains (Lane 1: Size Markers; 2: WT; 3: Mutant Δ*TRG1*)

**Table 2.**
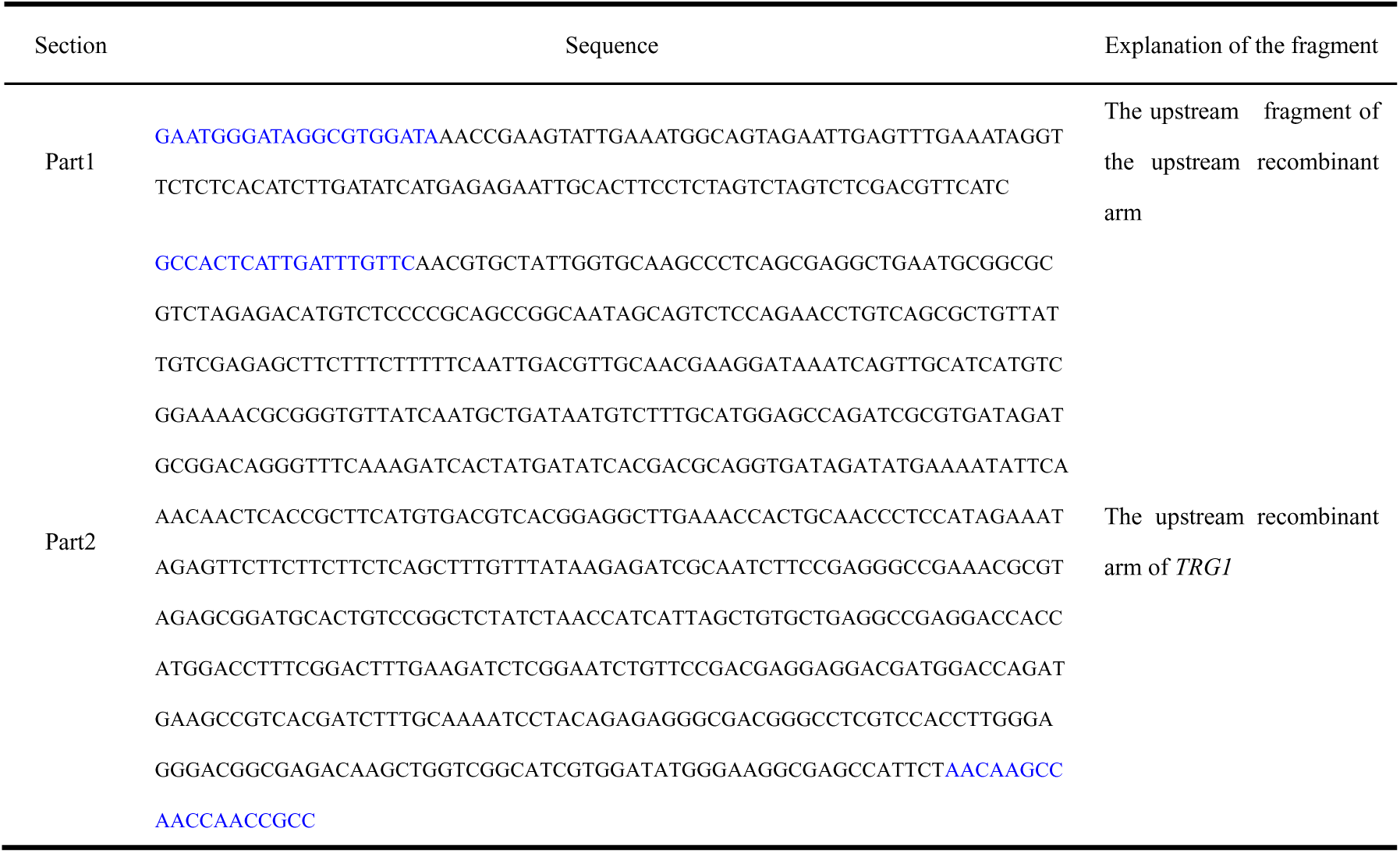
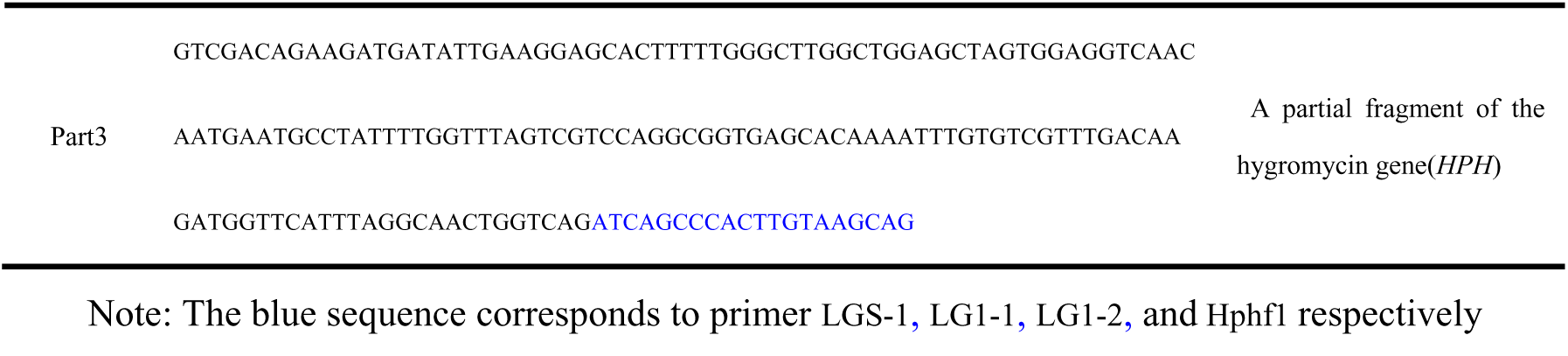
The detected recombinant sequence.

Total RNA was extracted from both WT and mutant *ΔTRG1*, reversed transcribed, and the resulting cDNA was used as a template for PCR amplification of the *TRG1* gene using the primer pair LG-Q1/LG-Q2. The results confirmed that the *TRG1* gene was normally transcribed in the WT strain, but TRG1 transcription could not be be amplified from the Δ*TRG1* mutant(Fig. 3C).

### 3.3 The mutant *ΔTRG1* showed no significant difference in morphology, growth rate and spore production compared to the WT

Phenotypic characteristics of *T. longibrachiatum* SMF2 (WT) and the mutant *ΔTRG1* on PDA medium. Observation, colony diameter measurement, and calculation of the relative growth rate showed no significant difference in growth ability between the two strains within 7 days of culture (Fig. 4).

**Figure 4.**
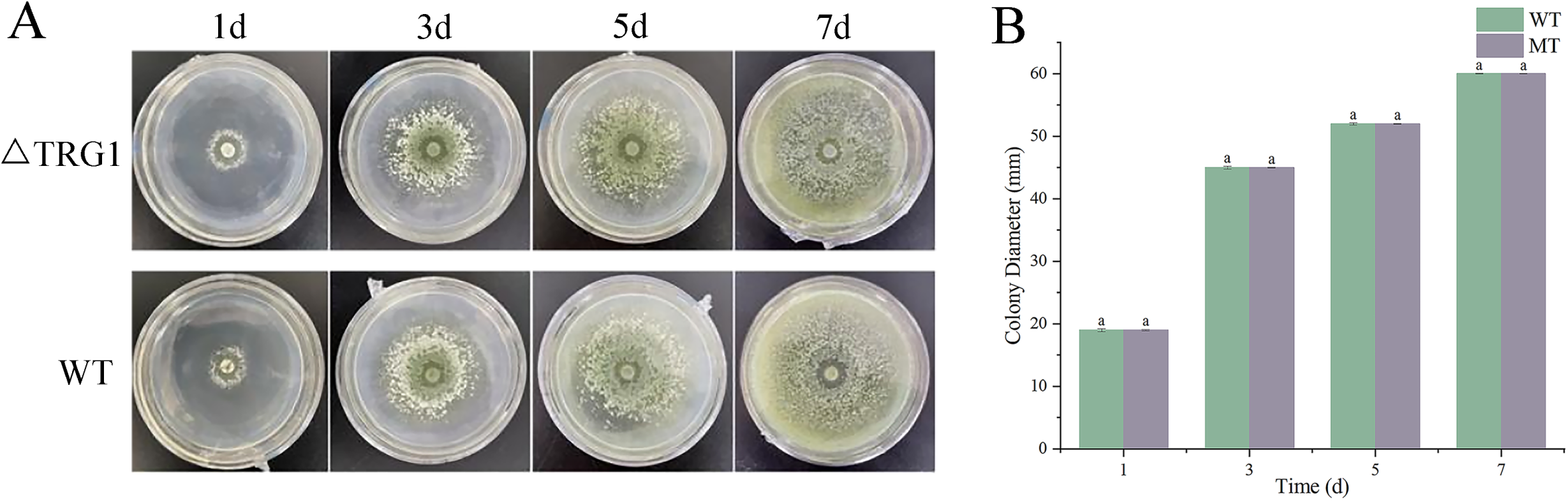
Comparison of growth rates between mutant *ΔTRG1* and the wild-type. Visual observations of colony growth over 7 days of culture (WT*: T. longibrachiatum* SMF2; *ΔTRG1:TRG1* knockout mutant). (A). Measurement of colony diameter of the WT and *ΔTRG1.* Data represent the mean ± sd. (n = 3). Different letters above the bars indicate a significant difference between the two strains within each timepoint as determined by an LSD test (p < 0.05).

Microscopic observation of the hyphae and spores revealed that both strains began to produce septa3 days after spore germination (Fig. 5). Both exhibited conidiophores branching straight to both sides, with spores single or clustered at the top. While the conidiophores of the mutants were generally shorter than those of the wild type (Fig. 5), this was a subtle difference.

**Figure 5.**
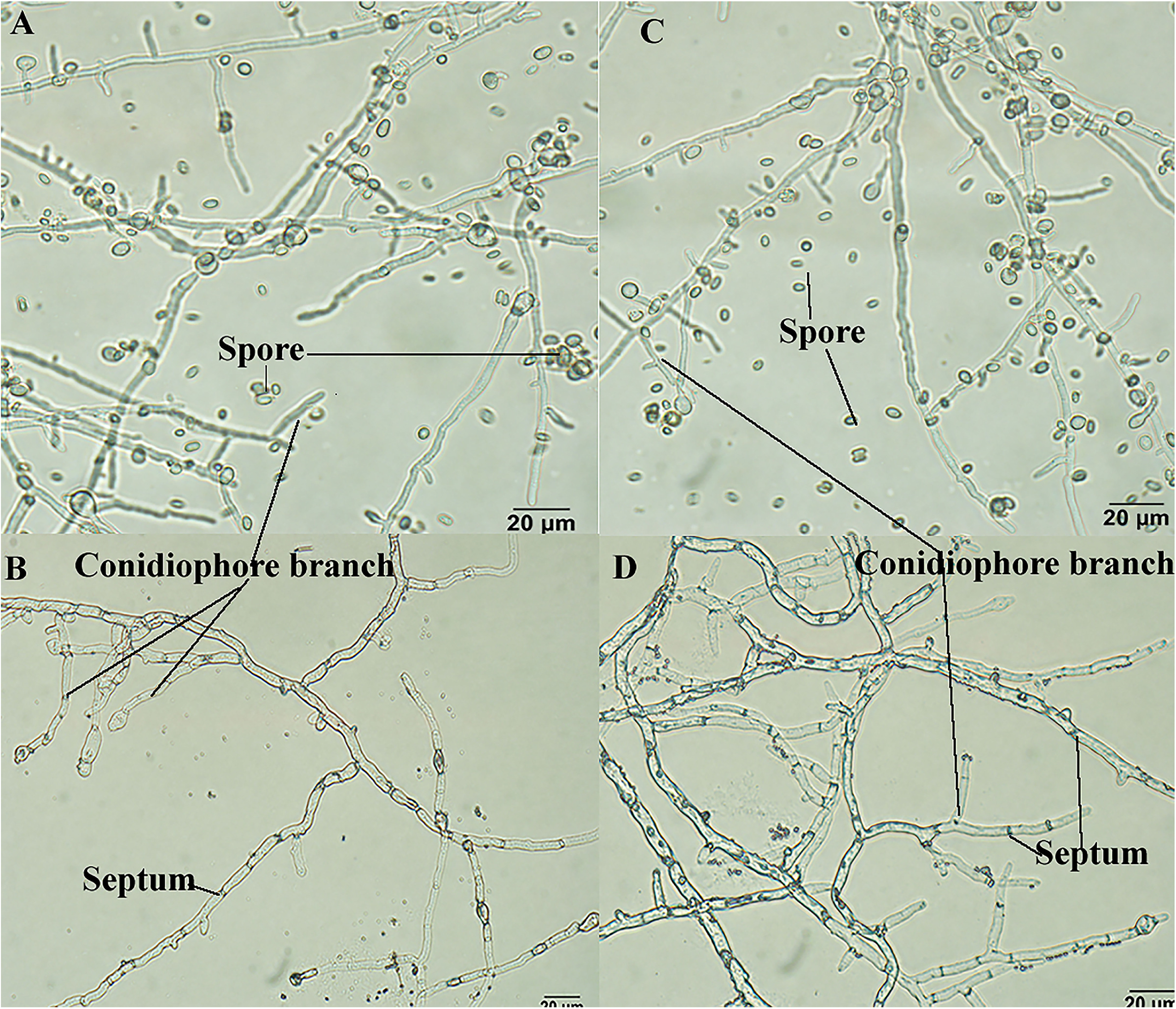
Comparison of hyphae and spores between mutant *ΔTRG1* and the wild-type. A/B: mutant *ΔTRG1*; C/D: the wild-type

Preparation of pore suspensions and hemocytometer counting showed that both *ΔTRG1* and WT spores were oval or nearly elliptical.The spore size of WT was (2.75∼4.21) μm × (5.60∼6.92) μm (width by length), while the spore size of the *ΔTRG1* was (2.67∼4.05) μm × (4.43∼5.95) μm, showing no significant change in spore size or yield (Fig. 5). These results indicate that the disruption of the TRG1 gene does not significantly affect the primary morphological or reproductive characters of the fungus (Fig. 5).

### 3.4 The mutant Δ*TRG1* showed a significantly lower inhibitory effect on *Botrytis cinerea*

The inhibition activity of *T. longibrachiatum* SMF2 (WT) and the mutant *ΔTRG1* against *B. cinerea* was assessed in vitro (Fig. 6). Both strains were able to inhibit the growth of *B. cinerea* over the 4-day culture period. On the fourth day, the inhibition rates were 71% for the WT and 60% for the mutant Δ*TRG1* (Table 3), with the WT strain demonstrating a significantly higher inhibition rate than the mutant Δ*TRG1* (LSD test, (p < 0.05).

**Figure 6.**
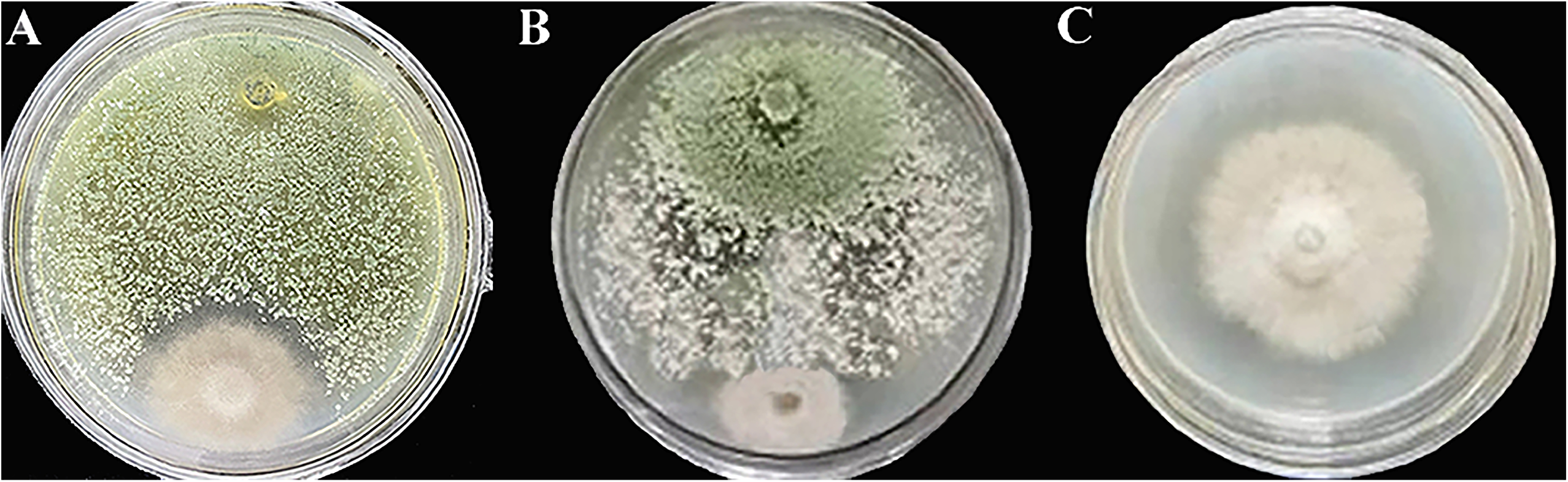
In vitro inhibition of *Botrytis cinerea* by WT and mutant ΔTRG1 strains when co-cultured on PDA. A: *ΔTRG1* and *B. cinerea*; B: WT and *B. cinerea* ; C: *B. cinerea*

**Table 3.**
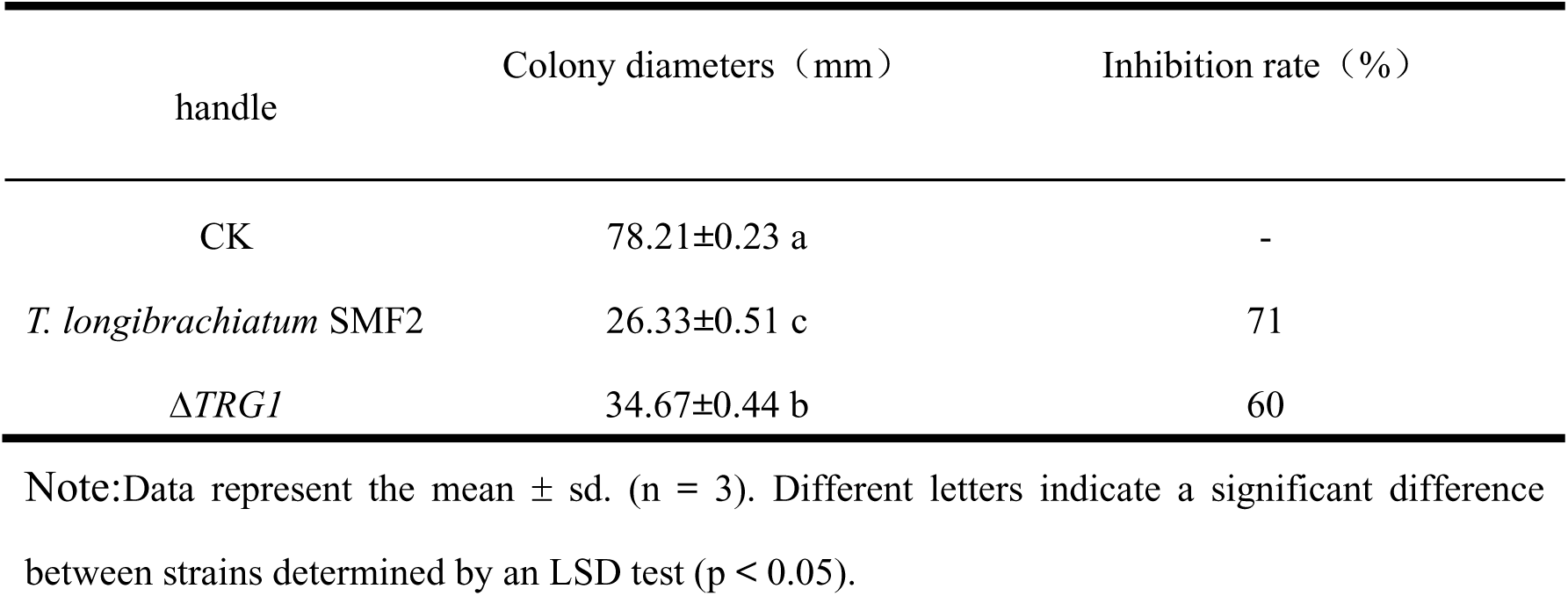
Inhibition rate of the WT and mutant *ΔTRG1* on *Botrytis cinerea*.

| handle | Colony diameters (mm) | Inhibition rate (%) |
| --- | --- | --- |
| CK | 78.21±0.23 a | - |
| *T. longibrachiatum* SMF2 | 26.33±0.51 c | 71 |
| $\Delta TRG1$ | 34.67±0.44 b | 60 |

Note:Data represent the mean ± sd. (n = 3). Different letters indicate a significant difference between strains determined by an LSD test ( $p < 0.05$ ).

### 3.5 The control effect of mutant *ΔTRG1* on *Botrytis cinerea* was reduced on detached rose leaves

After spraying the spore suspension of mutant Δ*TRG1* and the WT on Rose leaves both strains alleviated gray mold symptoms compared with the control group. However, the WT strain showed a generally higher ability to decrease disease symptoms than that of mutant Δ*TRG1* (Fig. 7A; Table 4).

**Figure 7.**
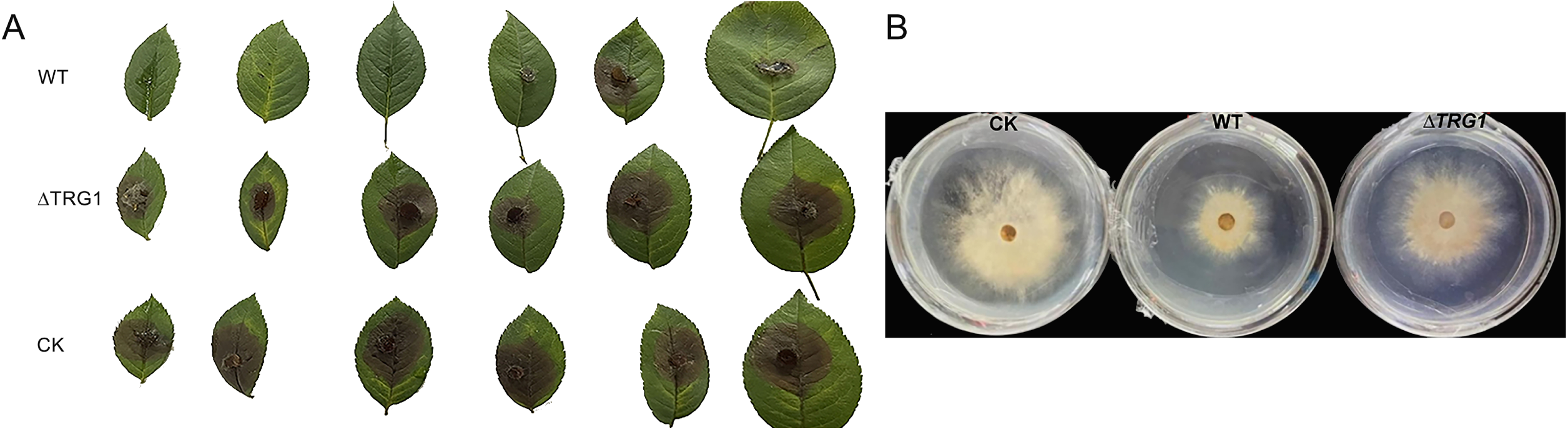
A: The protective effect of the fermentation broth of *ΔTRG1* mutant and the WT strains on the gray mold disease of detached rose leaves B: In vitro inhibitory effect of the fermentation broth of the *ΔTRG1* mutant and WT strains on *B. cinerea*. (CK:The Control ; WT: *T. longibrachiatum* SMF2 ; *ΔTRG1*: *TRG1* knockout mutant)

**Table 4.** The control effect of the WT and mutant Δ*TRG1* against rose gray mold disease of detached leaves and petals.

| handle | Leaves Disease index | Leaves protective effect | Petals Disease index | Petals protective effect |
| --- | --- | --- | --- | --- |
| CK | 95.83±2.12 a | - | 91.25±1.98 a | - |
| *T. Longibrachiatum*<br>SMF2 | 29.16±1.63 c | 69.8±3.49 a | 25.02±0.87 c | 72.58±3.49 a |
| $\Delta TRG1$ | 70.83±1.86 b | 26.08±2.58 b | 34.37±1.79 b | 62.33±2.58 b |

Note: Data represent the mean ± sd. (n = 3). Different letters indicate a significant difference between strains determined by an LSD test ( $p < 0.05$ ).

Seven days after treatment, , the disease index of the leaves treated with the mutant Δ*TRG1* was 70.83, which was significantly higher than that of the WT strain (29.16)). Correspondingly, the protective effect of Δ*TRG1* was 26.08%, which was significantly lower than the 69.8 % observed for the WT strain (LSD test, (p < 0.05) (Table 4).

### 3.6 The inhibitory effect of the fermentation broth of the mutant *ΔTRG1* on *B. cinerea* was lower than that of the WT

The cell free fermentation broths from both WT and *ΔTRG1* strains showed certain inhibitory effects on *B. cinerea*. However, the inhibitory effect of the Δ*TRG1* mutant broth was significantly lower than that of the WT strain, with an inhibition rate of 29% for the mutant Δ*TRG1* and 60% for the WT on the 4th day after inoculation (Fig. 7B; Table 5). This result strongly suggests that the reduced biocontrol ability of Δ*TRG1* is due to a reduction in secreted, active secondary metabolites.

**Table 5.** Inhibition rate of the WT and mutant Δ*TRG1* fermentation broth on *Botrytis cinerea*.

| Treatment | Colony diameters (mm) | Inhibition rate (%) |
| --- | --- | --- |
| CK | 78.21±0.23 a | - |
| *T. Longibrachiatum* SMF2 | 34.6±0.42 c | 60 |
| $\Delta TRG1$ | 57.16±0.21 b | 29 |

Note: Data represent the mean ± sd. (n = 3). Different letters indicate a significant difference between strains determined by an LSD test ( $p < 0.05$ ).

### 3.6 After the treatment of the mutant*ΔTRG1* fermentation broth, the activity level of the resistance-related enzymes in the detached rose leaves was lower than that of the WT

The effect of the spore-free fermentation broths on the induction of systemic resistance in rose leaves was assessed by measuring soluble protein, superoxide dismutase (SOD), and catalase (CAT) activities.

One day after treatment, the content of soluble protein was significantly higher in both treatment groups than in the control, with the WT treatment resulting in higher content than the *ΔTRG1* mutant. From the third day onward, the soluble protein of both treatments began to decrease (Fig. 8).

**Figure 8.**
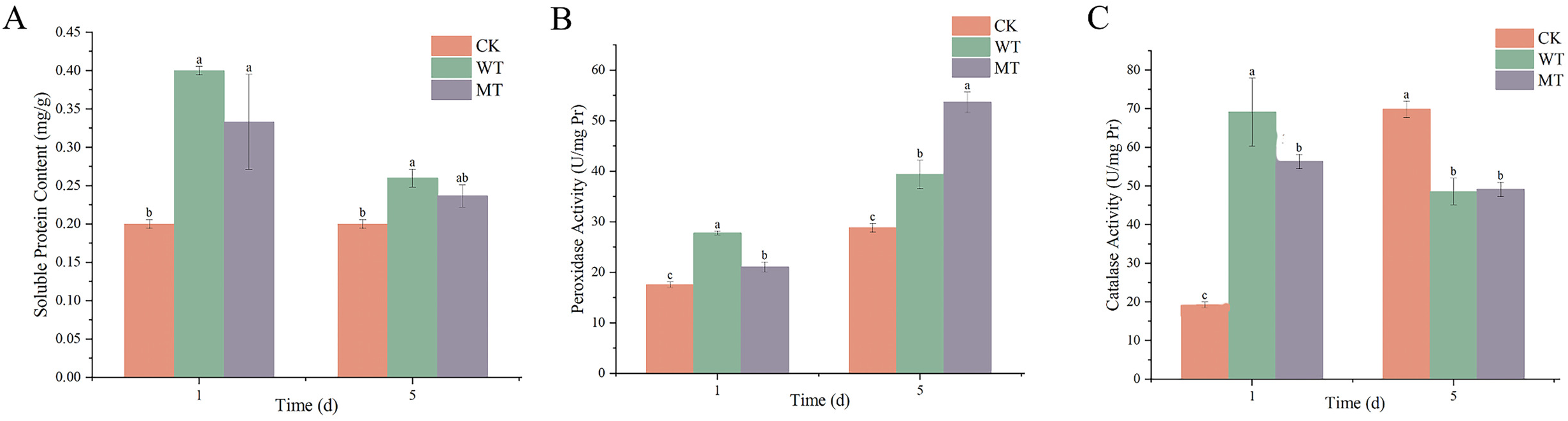
The activity level of the resistance-related enzymes indetached rose leaves after treatment with WT and Δ*TRG*1 mutant fermentation broth. A: soluble protein; B: POD activity; C: CAT activity

Similarly, the activities of SOD and CAT in rose leaves were higher than those of the control. Notably, the activities of both SOD and CAT in leaves treated with the WT fermentation broth were significantly higher than those treated with the mutant *ΔTRG1* on the 1st day (Fig. 8).

### 3.7 HPLC results showed that the yield of peptaibols produced by the Δ*TRG1* was significantly reduced

Analysis of the fermentation extracts by HPLC provided direct evidence of metabolite reduction (Fig. 9). Although both the wild-type strains and mutant Δ*TRG1* showed peptaibol peaks at 25 min, which corresponds to Trichokonin VI, the main active peptide of *T. longibrachiatum* SMF2), the difference in peak area size was significant. Peak area calculation showed that the peptaibol yield of *T. longibrachiatum* SMF2 (WT) was 2.5 times higher than that of the mutant Δ*TRG1*(Fig. 9). This result directly confirms that the transcription regulator gene TRG1 is essential for the high-yield production of peptaibols, explaining the observed reduction in all functional biocontrol assays.

**Figure 9.**
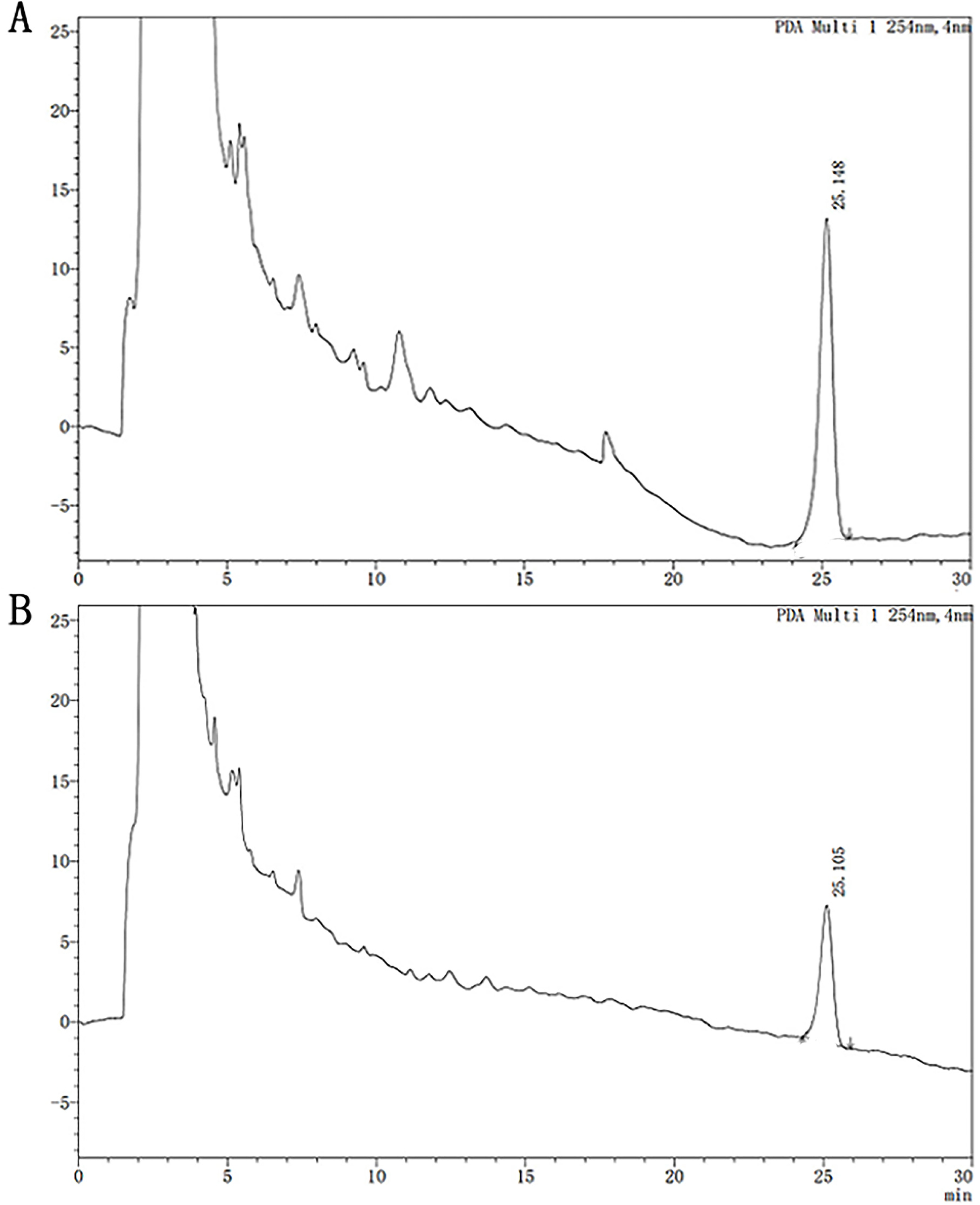
Comparise of the yield of peptaibols produced by the WT and mutant ΔTRG1. A:The WT; B: mutant Δ*TRG1*

## 4. Discussion

*Trichoderma spp*. are important biocontrol agents, effective against numerous pathogens including fungi (*Fusarium oxysporum*, *B. cinerea*), oomycetes (*Phytophthora infestans*), and bacteria (*Erwinia*) (María et al., 2023; Li et al., 2014). These fungi produce a variety of secondary metabolites that facilitate the inhibition of pathogenic microorganisms, promote plant growth, and enhance their own colonization(Jorge et al., 2019; Jacobus et al., 2020; Ravindra et al., 2020). Among these metabolites, peptaibols, synthesized through NRPS pathways, are particularly important for microbial inhibition, plant growth promotion, and the induction of plant resistance (Song et al., 2006).

Research has established that NRPS genes are massive, with some, like *tex1*, encoding proteins exceeding 20,000 amino acid residues (Wiest, et al. 2002). Targeted disruption of these core NPR genes, such as *tex1* or t*ex2* leads to the loss or partial loss of peptaibol production (Wiest et al., 2002; Chutrakul and Peberdy, 2005; Mukherjee et al., 2011). Furthermore, the genes involved in peptaibol biosynthesis, processing, and transport often form NRPS gene clusters that include associated regulatory genes (Zhang et al., 2014; Lu et al., 2017). For instance, in *T. longibrachiatum* SMF2, two NRPS genes, tlx1 and tlx2, responsible for the production of the 20-amino acid Trichokonin VI (TKA) and 12-amino acid TKB, have been identified (Xie et al., 2015; Zhou et al., 2019). Known regulatory elements, such as the glucose sensor homolog TlSTP1 and the LaeA homolog TlLAE1, have been shown to play important roles in regulating peptaibol production in this species (Shi et al., 2020). Despite this progress, many questions regarding the precise transcriptional regulatory mechanisms remain unresolved.

In this study, bioinformatics analysis confirmed that the predicted gene cluster containing the NRPS gene *tlx1* included genes that may be involved in peptide modification, transport, and synthesis regulation. We focused on the transcription regulator gene *TRG1* by constructing a knockout mutant (Δ*TRG1*). Phenotypic characterization showed that the Δ*TRG1* mutant exhibited no significant alterations in growth rate, colony morphology, spore production, and spore size compared to the WT strain, indicating TRG1 may not be essential for primary metabolism. However, functionally, the Δ*TRG1* mutant exhibited lower in vitro inhibition activity against *B. cinerea* and poorer in vivo control effect on gray mold compared. Moreover, the *ΔTRG1* cell-free fermentation broth was less effective at inducing resistance-related enzymes (SOD and CAT) in rose leaves. In particular, the HPLC analysis of the fermentation broth showed that the yield of peptaibols produced by mutant was only 40 % of the WT yield (a 2.5-fold reduction). The synthesis of these anti-fungal and resistance-inducing metabolites is mediated by NRPSs, leading us to conclude that the transcriptional regulator gene TRG1 is intimately involved in the positive regulation of peptaibol synthesis. The genes involved in fungal secondary metabolism generally exist in gene clusters regulated by complex, multi-level networks, with transcriptional regulation being the most critical (Lu et al., 2017). Transcriptional regulation in filamentous fungi often involves two classes of transcription factors (TFs): global TFs, which respond to external stimuli and regulate multiple biosynthetic gene clusters (BGC), and pathway-specific TFs, which mediate highly selective regulation, usually affecting only the BGC where they are located (Lyu, et al., 2020; Fox, et al., 2008).

The transcriptional regulator TRG1 obtained in our study is located directly on the NRPS gene cluster, suggesting it belongs to a pathway-specific positive transcription factor. The phenotype of the ΔTRG1 mutant showed a 60% reduction in peptaibol yield, strongly supporting this classification. This suggests that TRG1 acts as a powerful enhancer of transcription for relevant genes within the tlx1 cluster, potentially by binding to promoter regions. However, since peptaibol production did not completely cease, it implies that TRG1 is not an indispensable primary activator but rather a key positive modulator, suggesting that a basal level of expression or regulation by other factors remains.

Identifying transcriptional regulatory proteins like TRG1 is essential for understanding the molecular mechanisms underlying the synthesis of these valuable secondary metabolites. This knowledge is paramount for promoting target product yield and preparing highly efficient biocontrol strains through targeted genetic engineering. To fully elucidate the regulatory network, our next steps will involve further investigation: (1) Comparative transcriptome analysis and creation of double mutants to identify which global transcription factors regulate TRG1 expression; (2) Yeast one-hybrid assay to precisely locate the cis-elements bound by TRG1 on cluster promoters; and (3) Yeast two-hybrid analysis to explore potential interacting proteins. This comprehensive molecular approach will clarify the mechanism of TRG1 regulating peptaibols synthesis, providing the necessary molecular targets for rational bioengineering strategies to significantly improve peptaibol yields for agricultural applications.

## Acknowledgments

We thank Prof. Zhang Yuzhong in Shandong University for providing the strains and the genome sequence of *T. longibrachiatum* SMF2. This work was supported by Nature Science Fund of Shandong Province (ZR2020MC125); Liaocheng University-Enterprise Joint Project (K24LD171) and The Open Project of Liaocheng Universtiy Landscape Architecture Discipline (319462212).

## Author Contributions

Conceived of or designed study: Peibao Zhao, Aizhi Ren, Xiusheng Zhang; Performed research: Mengjiao Guan, Xiaoting Wang, Weishe Hu; Analyzed data: Hu Weishe, Wang Xiaoting, Li Ming; Writing: Hu Weishe, Aizhi Ren; Review and editing: Aizhi Ren, Peibao Zhao, Xiusheng Zhang; Funding acquisition: Peibao Zhao, Aizhi Ren. All authors read and approved the manuscript.

